# E-ChRPs: Engineered Chromatin Remodeling Proteins for Precise Nucleosome Positioning

**DOI:** 10.1101/480913

**Authors:** DA Donovan, JG Crandall, OGB Banks, ZD Jensvold, LE McKnight, JN McKnight

**Affiliations:** Institute of Molecular Biology, University of Oregon, Eugene OR 97403; Department of Biology, University of Oregon, Eugene OR 97403

## Abstract

Regulation of chromatin structure is essential for controlling the access of DNA to factors that require association with specific DNA sequences. The ability to alter chromatin organization in a targeted manner would provide a mechanism for directly manipulating DNA-dependent processes and should provide a means to study direct consequences of chromatin structural changes. Here we describe the development and validation of engineered chromatin remodeling proteins (E-ChRPs) for inducing programmable changes in nucleosome positioning by design. We demonstrate that E-ChRPs function both *in vivo* and *in vitro* to specifically reposition target nucleosomes and entire nucleosomal arrays, and possess the ability to evict native DNA-binding proteins through their action. E-ChRPs can be designed with a range of targeting modalities, including the SpyCatcher and dCas9 moieties, resulting in high versatility and enabling diverse future applications. Thus, engineered chromatin remodeling proteins represent a simple and robust means to probe regulation of DNA-dependent processes in different chromatin contexts.

## Introduction

The nucleosome is the fundamental repeating unit of chromatin, composed of DNA wrapped around an octamer of histone proteins. While nucleosomes are dynamic structures that are constantly assembled, disassembled, and repositioned in the genome, their positions at gene regulatory elements like transcription start sites (TSSs) show characteristic organization (Lai and Pugh, 2017). Thus, nucleosome positions are thought to have regulatory implications for DNA-dependent processes like transcription, replication and DNA repair (Hauer and Gasser, 2017; MacAlpine and Almouzni, 2013; Venkatesh and Workman, 2015). Because positions of nucleosomes in the genome play a major role in determining DNA sequence accessibility, the ability to precisely manipulate nucleosome positions would have profound implications for investigating and controlling DNA-dependent processes *in vivo*.

ATP-dependent chromatin remodeling factors couple the hydrolysis of ATP to the movement of nucleosomes along a fragment of DNA (Cairns et al., 1996; Fazzio and Tsukiyama, 2003; Langst et al., 1999; Smith and Peterson, 2005; Stockdale et al., 2006; Tsukiyama et al., 1994). By altering the positions of nucleosomes, this family of enzymes controls the accessibility of underlying DNA *in vivo*, thereby regulating DNA-dependent processes. The CHD and ISWI families of chromatin remodelers contain a conserved catalytic ATPase that drives chromatin remodeling by binding and hydrolyzing ATP (Zhou et al., 2016), and a C-terminal region that interacts with extranucleosomal DNA to modify the direction and outcome of nucleosome repositioning (Gangaraju and Bartholomew, 2007; Hota et al., 2013; McKnight et al., 2011; Ryan et al., 2011).

Previous work established that chromatin remodeling by *S. cerevisiae* Chd1 can be targeted to specific nucleosomes by replacing the native, nonspecific Chd1 DNA binding domain (DBD) with sequence-specific DBDs (McKnight et al., 2011; Nodelman and Bowman, 2013). We previously showed that hybrid Chd1 fusions with exogenous, sequence-specific DBDs predictably move nucleosomes onto their recruitment sequences *in vitro* (McKnight et al., 2011). We recently demonstrated that fusion of Chd1 to the Zn2Cys6 DBD from Ume6, a meiotic repressor from yeast, allows directed nucleosome positioning at target genes across the *S. cerevisiae* genome (McKnight et al., 2016).

Here we have simplified and greatly expanded the customizable design and validated the function of sequence-targeted chromatin remodeling proteins using diverse targeting strategies. These <u>e</u>ngineered <u>ch</u>romatin <u>r</u>emodeling <u>p</u>roteins (E-ChRPs) work with a wide variety of targeting domains and can occlude target DNA sequences by precisely repositioning nucleosomes onto recruitment motifs. We show that E-ChRPs possessing transcription factor (TF) DNA binding domains can block binding of endogenous transcription factors by incorporating TF binding sites into nucleosomes genome-wide. E-ChRPs can also be directly recruited to DNA-associated TFs through SpyTag/SpyCatcher pairs (Zakeri et al., 2012) allowing for identification and occlusion of TF-bound genomic loci. Finally, we show that positioning of nucleosomes can be achieved by a dCas9-targeted E-ChRP using optimized, noncanonical gRNAs.

## Design

The core E-ChRP design was inspired by previous work (McKnight et al., 2011; McKnight et al., 2016) where individual sequence-specific DNA binding domains (DBDs) replaced the C-terminal nonspecific DNA binding domain of a functional *S. cerevisiae* Chd1 chromatin remodeler fragment (Fig 1A). Yeast Chd1 is an ideal enzyme for engineered chromatin remodeling because it is monomeric, displays robust nucleosome positioning activity on nucleosome substrates derived from multiple organisms, and is less influenced by histone modifications than other chromatin remodelers (Ferreira et al., 2007; Hauk et al., 2010). After the Chd1 catalytic module we incorporated unique restriction sites flanking the targeting domain in vectors allowing for recombinant expression in *E. coli*, constitutive expression from ADH1 or GPD promoters in *S. cerevisiae* (Mumberg et al., 1995), or galactose-inducible expression from the HO locus in *S. cerevisiae* (Voth et al., 2001). This scaffold allows easy swapping of the C-terminal targeting domain resulting in a simple method to design chromatin remodelers that can be localized to desired nucleosomes. To demonstrate the versatility of the approach, we incorporated and assessed engineered chromatin remodeling through multiple transcription factor DNA binding domains, through SpyCatcher/SpyTag pairs, and through dCas9 targeting (Fig 1B). We first assessed the ability of different E-ChRPs to reposition target-containing mononucleosomes in a purified biochemical assay (Eberharter et al., 2004). To validate *in vivo* function, we introduced E-ChRPs into *S. cerevisiae* and measured global nucleosome positions by MNase-seq. Functional E-ChRPs can position targeted nucleosomes onto recruitment motifs as measured by mononucleosome sliding toward recruitment sequences *in vitro* or target motif occlusion by nucleosomes *in vivo* (Fig 1C).

**Figure 1.**
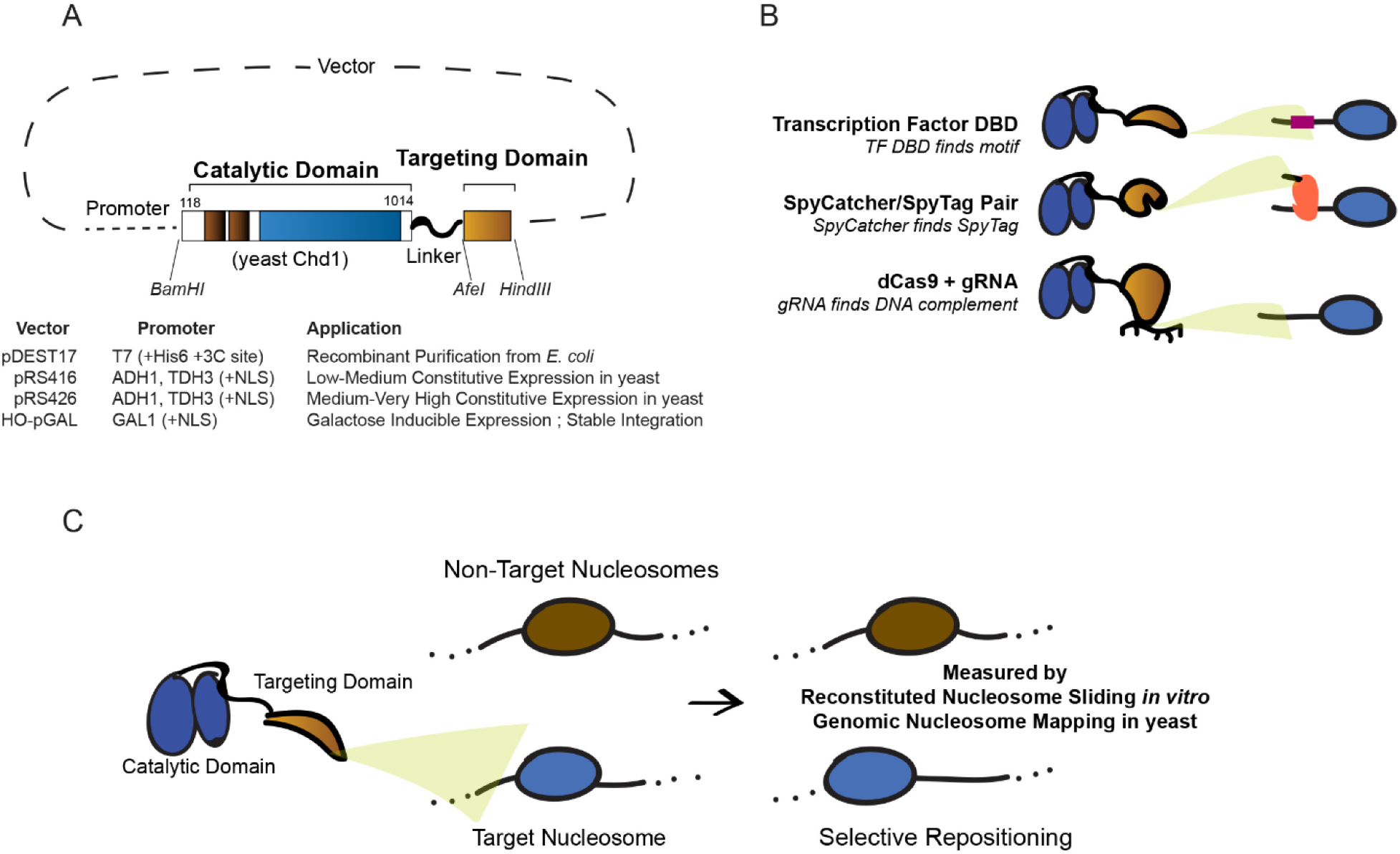
Strategies for Targeted Nucleosome Positioning by E-ChRPs. (A) General architecture of the E-ChRP core where the yeast Chd1 catalytic domain is linked to a targeting domain with a flexible linker. (B) Summary of targeting methods employed in this work, including: sequence-specific DNA binding domain targeting to a recognition motif (top), SpyCatcher domain covalently attaching to a SpyTag-containing chromatin-bound protein (middle), and dCas9-bound gRNA interacting with a complementary sequence (bottom). (C) Predicted outcome from targeted E-ChRPs, indicating select nucleosomes are positioned by the E-ChRP onto the recruitment site.

## Development and Optimization of a Targeted Remodeler Core

We previously demonstrated that fusion of a foreign DNA binding domain to the Chd1 catalytic core leads to occlusion of a recruitment motif by targeted and directional repositioning of nucleosomes (McKnight et al., 2011; McKnight et al., 2016). Though functional both *in vitro* and *in vivo*, remodeler fusions where the DNA binding domain was directly fused to the Chd1 core resulted in a limited “reach” and nucleosomes residing further than 20 base pairs from the DNA recognition element were not efficiently moved. In addition, the creation of new remodeler fusion proteins was previously cumbersome and lacked versatility. To address these limitations we first created the E-ChRP scaffold (Fig 1A), which consists of the catalytic core of the yeast Chd1 protein followed by a flexible linker including 11 repeats of the glycine-glycine-serine sequence that was previously shown to extend the Chd1 reach (Nodelman and Bowman, 2013). We next created an array of plasmids for recombinant bacterial expression or yeast constitutive or inducible expression allowing for one-step cloning of a desired fusion domain (Fig 1A).

We examined whether the addition of a flexible linker between the Chd1 remodeler core and DNA binding domain increased the reach of these E-ChRPs. We tested the ability of an E-ChRP with a DBD from the *S. cerevisiae* meiotic repressor Ume6 to move mononucleosomes containing a recognition motif, URS1 (Park et al., 1992), 20 or 40 base pairs from the nucleosome edge (Fig 2A). Without a flexible linker, the Chd1-Ume6 E-ChRP was active exclusively when the motif was 20bp away (compare lanes 1 with 3, 2 with 4). In contrast, addition of eleven repeats of glycine-glycine-serine (GGSx11) allowed the remodeler to efficiently mobilize both nucleosome substrates (compare lanes 1 with 5, 2 with 6). Additionally, the final location of the positioned nucleosomes was dependent on the location of the recognition motif (Fig 2A, compare lanes 5 and 6). Consistent with this increased reach *in vitro*, the Ume6 E-ChRP containing a GGSx11 linker positioned a larger fraction of distal nucleosomes onto target sequences across the *S. cerevisiae* genome (Fig 2B, C). Because the flexible linker led to more robust E-ChRP activity and our design was compatible *in vitro* and *in vivo*, we employed this general scaffold in all subsequent experiments.

**Figure 2.**
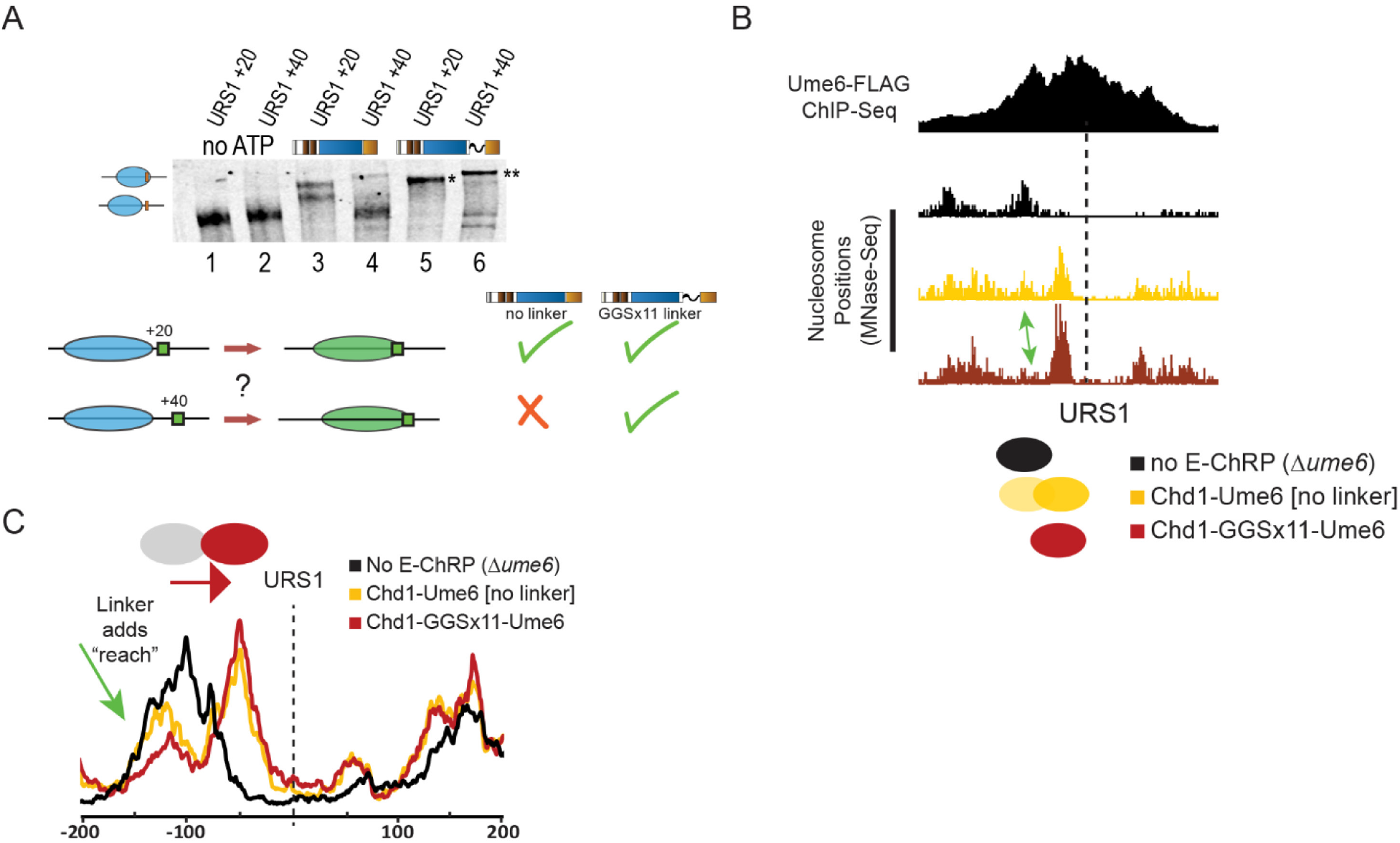
A Flexible Linker Increases the Reach of E-ChRPs *in vitro* and *in vivo*. (A) Nucleosome repositioning *in vitro* by Chd1-Ume6 with and without 11 repeats of glycine-glycine-serine between the Chd1 catalytic domain and Ume6 DNA binding domain. Nucleosomes with the Ume6 recognition motif (URS1) located 20 or 40 base pairs from the nucleosome edge were incubated with Chd1-Ume6(DBD) or Chd1-GGSx11-Ume6(DBD) and nucleosome were resolved by native PAGE (top). Nucleosome positions before and after remodeling were resolved by 6% native PAGE. Summary of repositioning of each substrate by Chd1 fusions (bottom). (B) Genome Browser image showing nucleosome positions at a representative Ume6 binding site for a parental strain lacking endogenous Ume6 (black), after introduction of Chd1-Ume6 without (yellow) or with (red) a flexible linker. Dashed line indicates the location of the Ume6 binding motif. Arrow indicates nucleosome that is more efficiently positioned with a Chd1-GGSx11-Ume6(DBD) fusion. (C) Average nucleosome positioning at genomic Ume6 binding sites showing Chd1-GGSx11-Ume6(DBD) more readily positions nucleosomes distal to the recognition element than Chd1-Ume6(DBD) lacking a flexible linker.

## Remodeler Fusions are Highly Versatile *in vitro* and *in vivo*

We next tested mononucleosome targeting of multiple E-ChRPs with various DNA binding domains. We fused the DBD from *E. coli* AraC, *S. pombe* Res1, *D. melanogaster* Engrailed, or *R. norvegicus* Glucocorticoid Receptor to the E-ChRP scaffold. To determine if these E-ChRPs were functional on target nucleosomes *in vitro*, we generated end-positioned mononucleosomes assembled on the 601-positioning sequence (Lowary and Widom, 1998) with 125 base pairs of flanking DNA. The extranucleosomal DNA either contained or lacked a consensus binding motif corresponding to each different fusion tested. E-ChRPs containing DNA binding domains were able to mobilize nucleosomes containing their well-defined recruitment motifs (Ades and Sauer, 1994; Alroy and Freedman, 1992; Ayte et al., 1995; Niland et al., 1996) as measured by a native PAGE nucleosome sliding assay (Fig 3A). These E-ChRPs were inactive on nucleosomes lacking their respective motifs (Fig. 3A, lanes 22-27), demonstrating specificity for target substrates *in vitro*. Fusion of the native, sequence-nonspecific DBD from Chd1 to our E-ChRP scaffold showed no apparent discrimination against DNA sequences and was capable of fully mobilizing the nonspecific mononucleosome control (Fig. 3A, lanes 28-30).

**Figure 3.**
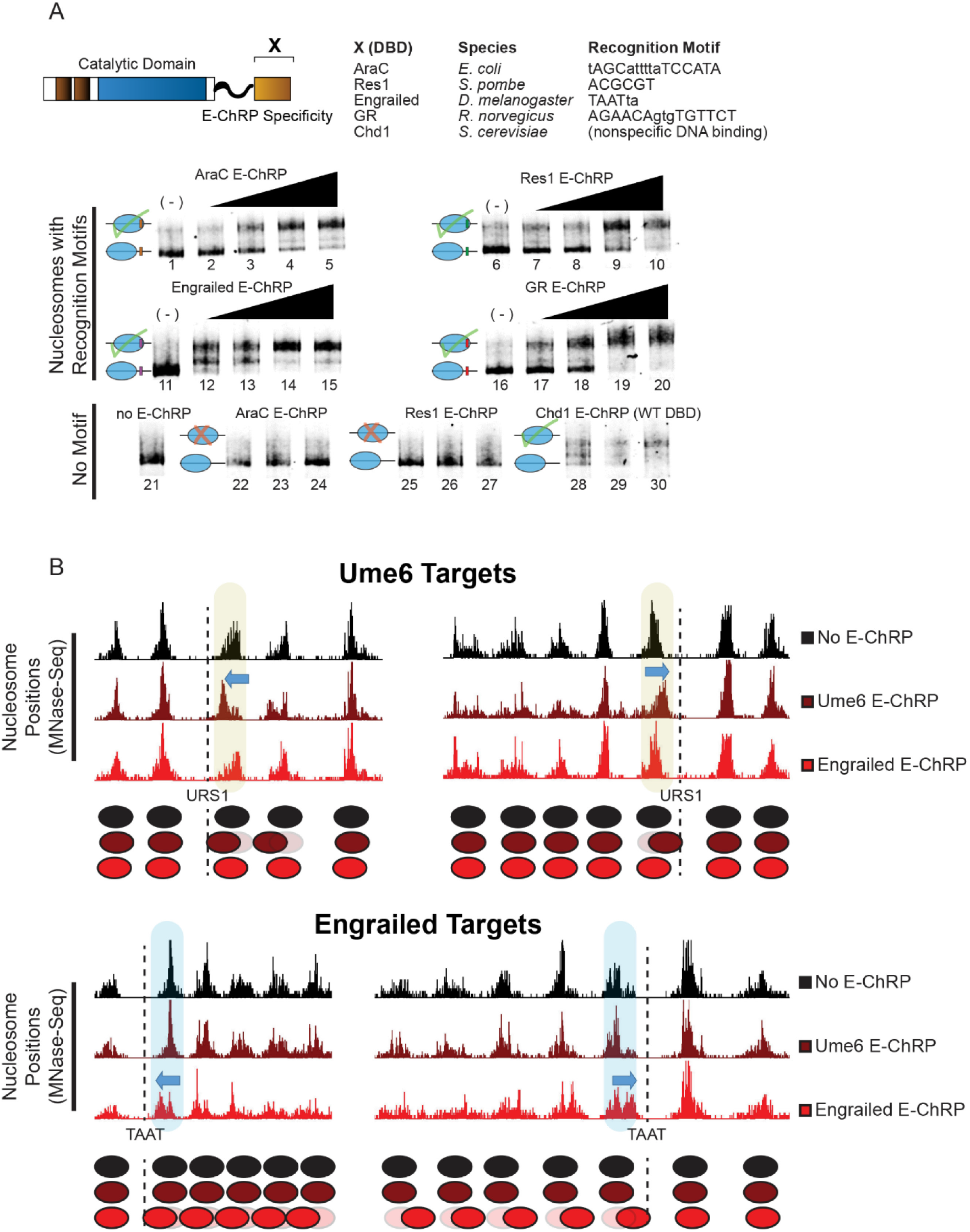
E-ChRPs with Distinct Transcription Factor DNA Binding Domains Specifically Position Target Nucleosomes *in vitro* and *in vivo*. (A) Nucleosome sliding assay demonstrating functionality of increasing concentrations of E-ChRPs containing AraC DBD (lanes 1-5, 22-24), Res1 DBD (lanes 6-10, 25-27), Engrailed DBD (lanes 11-15), Glucocorticoid Receptor DBD (lanes 16-20) or Chd1 endogenous DBD (lanes 28-30) in vitro. Nucleosomes in lanes 1-20 possess recognition motifs in extranucleosomal DNA for the respective E-ChRP (Ades and Sauer, 1994; Alroy and Freedman, 1992; Anderson et al., 1995; Ayte et al., 1995; Khan et al., 2018; Niland et al., 1996) while lanes 21-30 have no recognition motif. Lower electrophoretic mobility indicates repositioning of nucleosomes away from their end positions. (B) Yeast genomic nucleosome dyad positions are shown at representative Ume6 targets (URS1, top) or Engrailed targets (TAAT, bottom) in the presence or absence of Ume6 E-ChRP or Engrailed E-ChRP. Motif-proximal nucleosomes are highlighted next to indicated motifs, with blue arrows showing direction of nucleosome movement. Cartoon representations of nucleosome positions are provided for each locus. (See also Figure S1)

To determine whether E-ChRPs can be differentially targeted to specific subsets of nucleosomes *in vivo*, we introduced E-ChRPs into *S. cerevisiae* on a constitutive, ADH1-driven expression plasmid. When the E-ChRP contained a Ume6 DNA binding domain, nucleosomes were repositioned toward Ume6 binding motifs across the genome, but no nucleosome changes were detected at other genomic loci. Similarly, an E-ChRP containing the Engrailed DNA binding domain moved nucleosomes onto Engrailed motifs in the yeast genome without altering nucleosome positions at Ume6 binding motifs (Fig 3B). We also introduced Ume6 and Engrailed E-ChRPs into yeast under the high-expression GPD (TDH3) promoter on a 2μm plasmid (Mumberg et al., 1995). Expression of the Ume6 E-ChRP from this construct resulted in positioned nucleosomes at target sites without identified off-target activity similar to an ADH1-driven E-ChRP (Fig S1A). However, introduction of this higher expression plasmid containing an Engrailed E-ChRP only produced viable transformants in which the E-ChRP construct was deleted, truncated or mutated. This obligate inactivation of the Engrailed E-ChRP at high expression levels may result from promiscuous action of the Engrailed E-ChRP at tens of thousands of potential target sequences, which would presumably disrupt global nucleosome positioning in a deleterious and pleiotropic manner. Importantly, even when driven from the GPD (TDH3) promoter neither the Ume6 E-ChRP nor the Engrailed E-ChRP was active at target sites when the Chd1 remodeler core contained a catalytically-inactive Walker B (D513N) substitution (Hauk et al., 2010; Walker et al., 1982) (Fig S1A,B). Taken together, these results suggest that E-ChRPs can be specifically targeted through multiple distinct DNA binding domains that recognize sequence motifs with high or low complexity both *in vitro* and *in vivo*.

## E-ChRPs can Inducibly Remove Transcription Factors

To gain temporal control of E-ChRPs *in vivo*, we introduced the Ume6 E-ChRP under the control of a galactose-inducible promoter integrated at the HO locus in yeast (Voth et al., 2001). Prior to addition of galactose, endogenous Ume6 associates with its consensus sequence across the genome and cooperates with the ISW2 complex to position motif-proximal nucleosomes, leaving 30 bp between the nucleosome edge and URS1 motif (Goldmark et al., 2000; McKnight et al., 2016). After galactose induction of the Ume6 E-ChRP, a majority of nucleosomes nearest the URS1 site are efficiently repositioned to occlude the URS1 motif within two hours (Fig 4A). This galactose-inducible approach allows for more complete remodeling of nucleosomes than the same E-ChRP under the control of a constitutively active ADH1 promoter, potentially commensurate with differing expression levels under these distinct promoters (Fig 4A).

**Figure 4.**
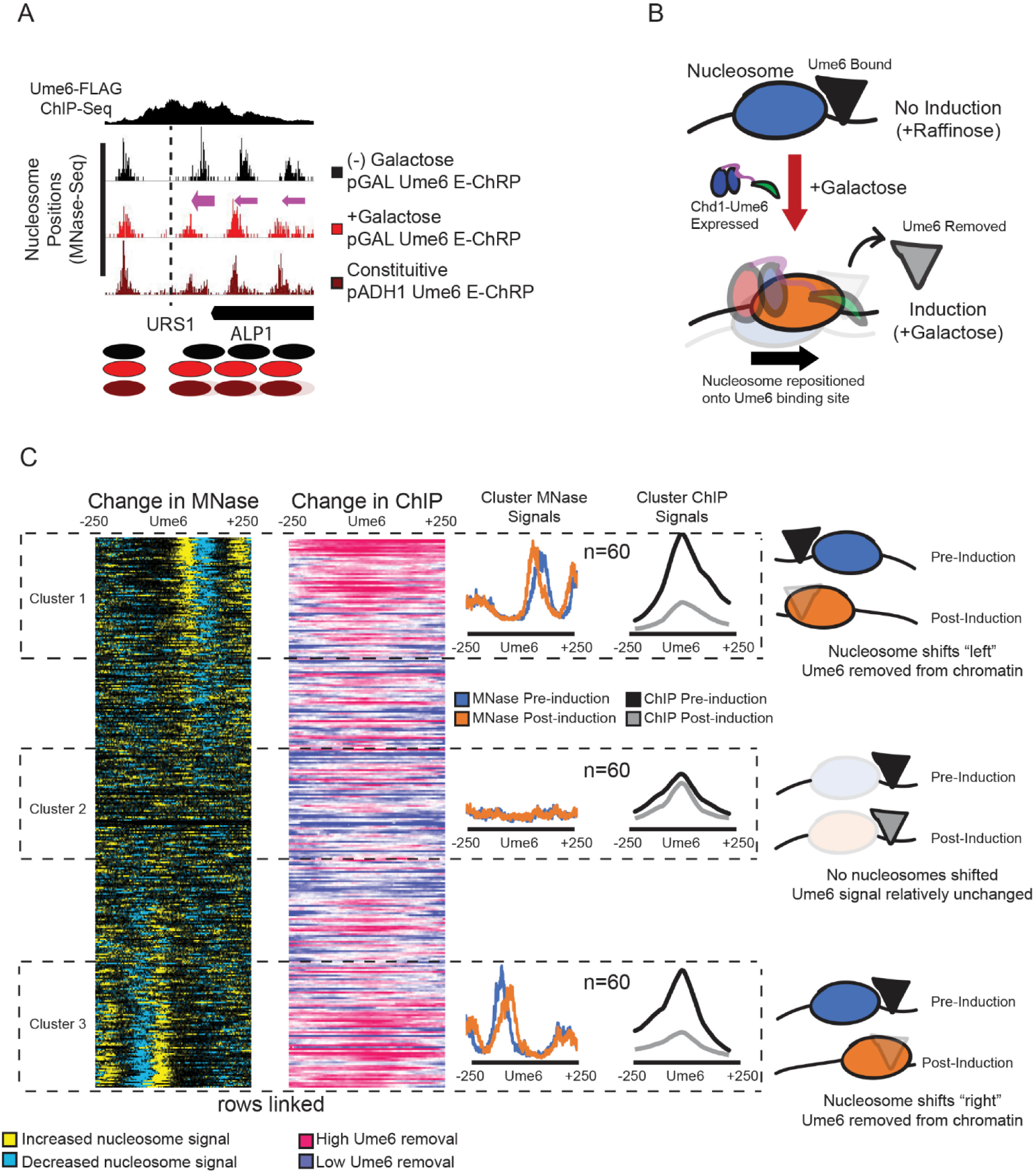
E-ChRPs can Inducibly Remove Endogenous Ume6 from Chromatin. (A) Nucleosome signal at a representative locus in yeast demonstrating positioning of nucleosomes toward recruitment motif (dashed line) by galactose-inducible Ume6 E-ChRP or a constitutively expressed E-ChRP under the ADH1 promoter. (B) Cartoon depiction of Ume6 E-ChRP activity blocking association of endogenous Ume6 at target sites. (C) Global analysis centered on 600 Ume6-FLAG ChIP peaks for change in nucleosome dyad signal (yellow/blue) after Ume6 E-ChRP induction and associated change in endogenous Ume6-FLAG ChIP signal (blue/red). Clusters represent Ume6 peaks with the highest and lowest change in nucleosome positioning based on ranked changes in MNase signal. Average nucleosome dyad signal for each cluster before and after E-ChRP induction is given in blue and orange, respectively. Change in Ume6-FLAG ChIP signal for each cluster is provided in black and gray traces, respectively. Cartoon interpretation for each cluster is provided on the right. (See also Figure S2)

We reasoned that because the post-induction nucleosome position results in the Ume6 recruitment motif becoming buried within nucleosomal DNA, remodeling by the Ume6 E-ChRP should interfere with binding of endogenous Ume6 (Fig 4B). To test this possibility, we tagged endogenous Ume6 with a FLAG epitope and measured Ume6-FLAG binding by ChIP-seq before and after induction of the Ume6 E-ChRP. Prior to induction, reproducible Ume6-FLAG binding was observed at URS1 sites across the genome (Fig S2A). After induction of the Ume6 E-ChRP, which shifted nucleosomes over URS1 sites, Ume6 binding (as measured by Ume6-FLAG ChIP signal) was strongly reduced or eliminated at many genomic locations (Fig S2A,B). When we sorted Ume6 binding sites based on whether proximal nucleosome positions were shifted after Ume6 E-ChRP induction, we noticed that loss of Ume6-FLAG signal was strikingly reduced where nucleosomes were shifted, but minimally reduced where nucleosomes were not shifted (Fig 4C, S2B). To verify that this reduction in Ume6-FLAG signal was not due to direct binding competition between endogenous Ume6-FLAG and the E-ChRP Ume6 DNA binding domain, we measured Ume6-FLAG ChIP signal in the presence of a catalytically inactive Ume6 E-ChRP. This construct, which cannot move nucleosomes but retains the Ume6 DNA binding domain, did not similarly reduce Ume6-FLAG signal (Fig S2A,B). Thus, E-ChRPs can inducibly move nucleosomes over target sequences to restrict access of the underlying DNA to endogenous DNA binding factors.

## SpyCatcher E-ChRPs Allow Simple Targeting to Chromatin-Bound Loci

One limitation of the above-described E-ChRPs is their need to compete with endogenous factors for binding sites. To circumvent this problem, we created an E-ChRP where the SpyCatcher protein is fused in place of a DNA binding domain in the Chd1 E-ChRP scaffold. SpyCatcher specifically recognizes a short (∼1kDa) SpyTag epitope, forming an isopeptide linkage that allows covalent protein fusions to be created *in vitro* and *in vivo* (Zakeri et al., 2012). This fusion provides two major improvements to the E-ChRP system. First, by simply appending SpyTag to different chromatin-binding factors of interest, nucleosome positioning can be achieved by a single SpyCatcher E-ChRP without the need to design new DBD fusions (Fig 5). Second, by tagging a transcription factor at its endogenous locus, the protein becomes a targetable element for the SpyCatcher E-ChRP only when bound to chromatin (Fig 5A). This bypasses the requirement of a vacant DNA binding site to target a DBD-containing E-ChRP, allowing access to sequences in the genome that could otherwise be blocked by a stably-bound transcription factor. In sum, this strategy produces a single SpyCatcher E-ChRP that can be targeted to any chromatin-bound protein of interest in the genome by simple attachment of a short SpyTag.

**Figure 5.**
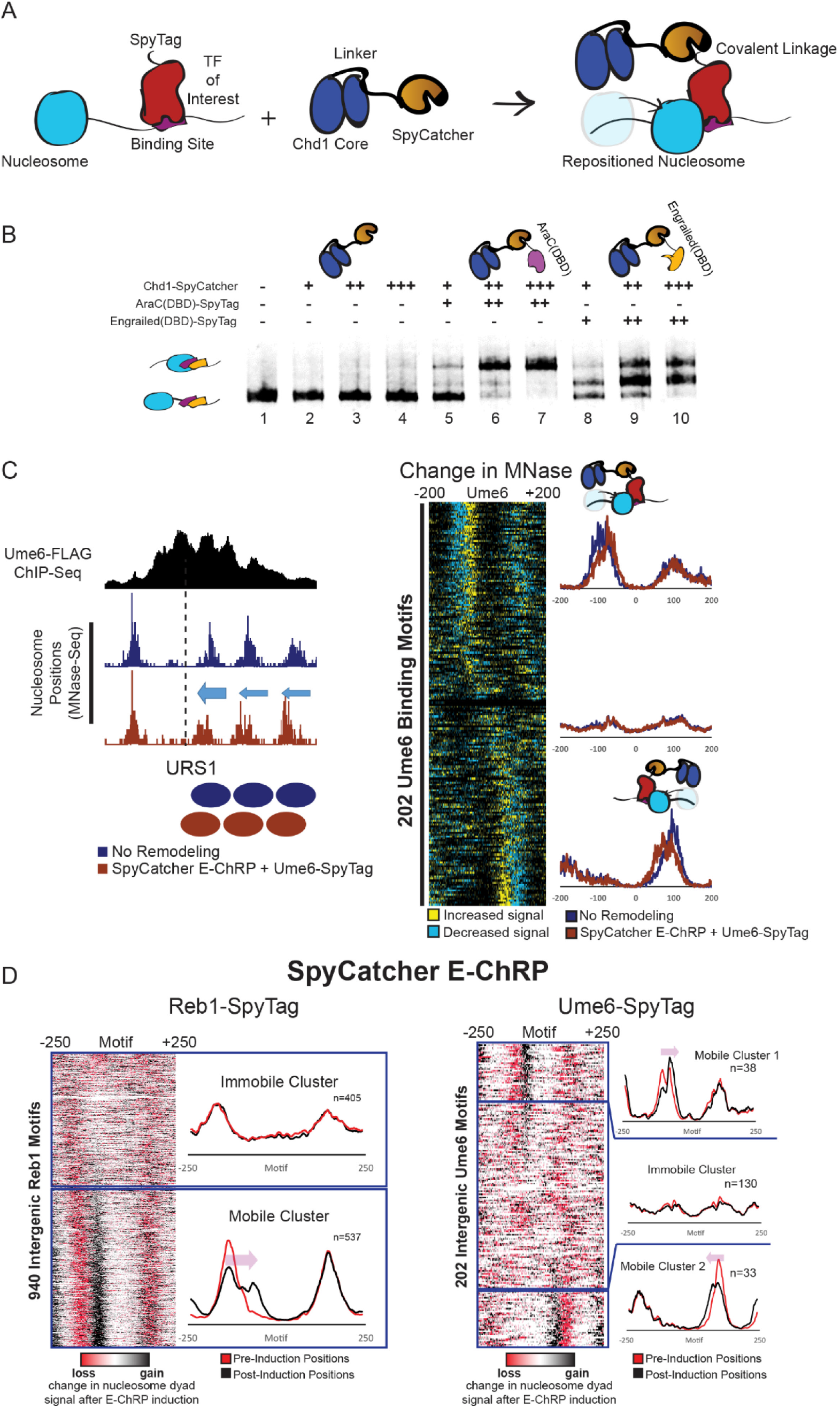
Development and Validation of E-ChRPs Containing SpyCatcher/SpyTag Pairs. (A) Cartoon representation for introducing a Chd1-SpyCatcher E-ChRP into cells containing SpyTagged, chromatin-bound proteins. The SpyCatcher domain forms a covalent isopeptide bond with SpyTag, allowing for localization of E-ChRP activity to endogenously-bound chromatin proteins. (B) Nucleosome sliding assay demonstrating that a single SpyCatcher E-ChRP cannot position nucleosomes without a SpyTag-containing DBD (lanes 2-4) but can use a SpyTagged AraC DBD (lanes 5-7) or Engrailed DBD (lanes 8-10) to reposition nucleosomes containing respective DBD recognition motifs. (C) Representative motif in yeast where ADH1-driven SpyCatcher E-ChRP can reposition nucleosomes at a Ume6 binding site in the presence of Ume6-SpyTag (left) and genomic analysis of nucleosome positioning by SpyCatcher E-ChRP at 202 intergenic instances of the Ume6 recognition sequence in cells containing SpyTagged Ume6 (right). (D) Genomic analysis of nucleosome positions in Reb1-SpyTagged cells (left) or Ume6-SpyTagged cells (right) before and after 2-hour induction of galactose-inducible SpyCatcher E-ChRP. Heat maps show change in nucleosome dyad signal after induction of SpyCatcher E-ChRP while individual traces show average positions of nucleosomes in each cluster before and after SpyCatcher E-ChRP induction for each SpyTag-DBD strain. (See also Figure S3)

To validate the function of the SpyCatcher E-ChRP design, we purified recombinantly-expressed Chd1-SpyCatcher and two SpyTag-containing DNA binding domains. Mononucleosomes harboring recognition sequences for each DNA binding domain in the extranucleosomal DNA were incubated with the SpyCatcher E-ChRP with and without addition of SpyTag-Engrailed(DBD) or SpyTag-AraC(DBD). The SpyCatcher E-ChRP has no activity on nucleosome substrates in the absence of a SpyTag-DBD pair (Fig 5B, lanes 2-4), since SpyCatcher has no intrinsic DNA binding affinity. However, addition of either SpyTag-AraC(DBD) (Fig 5B, lanes 5-7) or SpyTag-Engrailed(DBD) (Fig 5B, lanes 8-10) resulted in robust repositioning of mononucleosomes *in vitro*, demonstrating the versatility of this system. We next introduced the SpyCatcher E-ChRP under a constitutive ADH1 promoter into *S. cerevisiae* cells where a C-terminal SpyTag was added to full-length Ume6 at the endogenous locus. As expected, we observed repositioned nucleosomes at URS1 sites across the genome indicating chromatin remodeling at Ume6-bound loci (Fig 5C).

To achieve temporal control of this modular system *in vivo*, we appended SpyTag to the C-terminus of either Ume6 or Reb1, a yeast general regulatory factor, in a strain harboring a galactose-inducible SpyCatcher E-ChRP at the HO locus. After induction of SpyCatcher E-ChRP expression, nucleosomes were shifted toward Ume6 binding sites in cells containing Ume6-SpyTag or toward Reb1 binding sites in cells containing Reb1-SpyTag (Fig 5D, Fig S3A,B). Interestingly, the fraction of shifted nucleosomes was generally low at Ume6 binding sites in Ume6-SpyTag cells but comparatively higher at Reb1 binding sites in Reb1-SpyTag strains (Fig 5D). This difference could be explained by higher occupancy or stability of Reb1 than Ume6 binding at target sites, which would allow a greater fraction of Reb1-tethered SpyCatcher to mobilize motif-proximal nucleosomes. Consistent with this possibility, the cellular abundance of Ume6 is significantly lower than that of Reb1 (Kulak et al., 2014). For Reb1-SpyTag strains, the positioning of a single motif-proximal nucleosome by the SpyCatcher E-ChRP initiated the shift of an entire array of nucleosomes toward the target motif (Fig S3C), consistent with previous observations that the positioning of a “barrier nucleosome” influences and constrains positions of an entire array of nucleosomes (Mavrich et al., 2008; McKnight et al., 2016).

Interestingly, the positioning of nucleosomes appeared to occur on only the 5’ side of the Reb1 recognition sequence, suggesting the orientation of Reb1 binding impacts the ability of Chd1 to reach nucleosomes near binding sites (Fig 5D, Fig S3C). This restriction could be explained by a constrained C-terminus of Reb1 when bound to chromatin, which is consistent with similarly constrained Reb1-MNase cleavage patterns seen in previous ChEC-seq experiments (Zentner et al., 2015). Unexpectedly, the fraction of nucleosomes shifted at individual Reb1 binding sites varied greatly in our data set, with some sites exhibiting repositioning of nearly 100% of motif-proximal nucleosomes in the population and others having a much smaller fraction moved (Fig 6A-C). These differences are not explained by initial nucleosome occupancy or location differences (Fig 6B), but are possibly related to relative Reb1 occupancy at different genomic locations.

**Figure 6.**
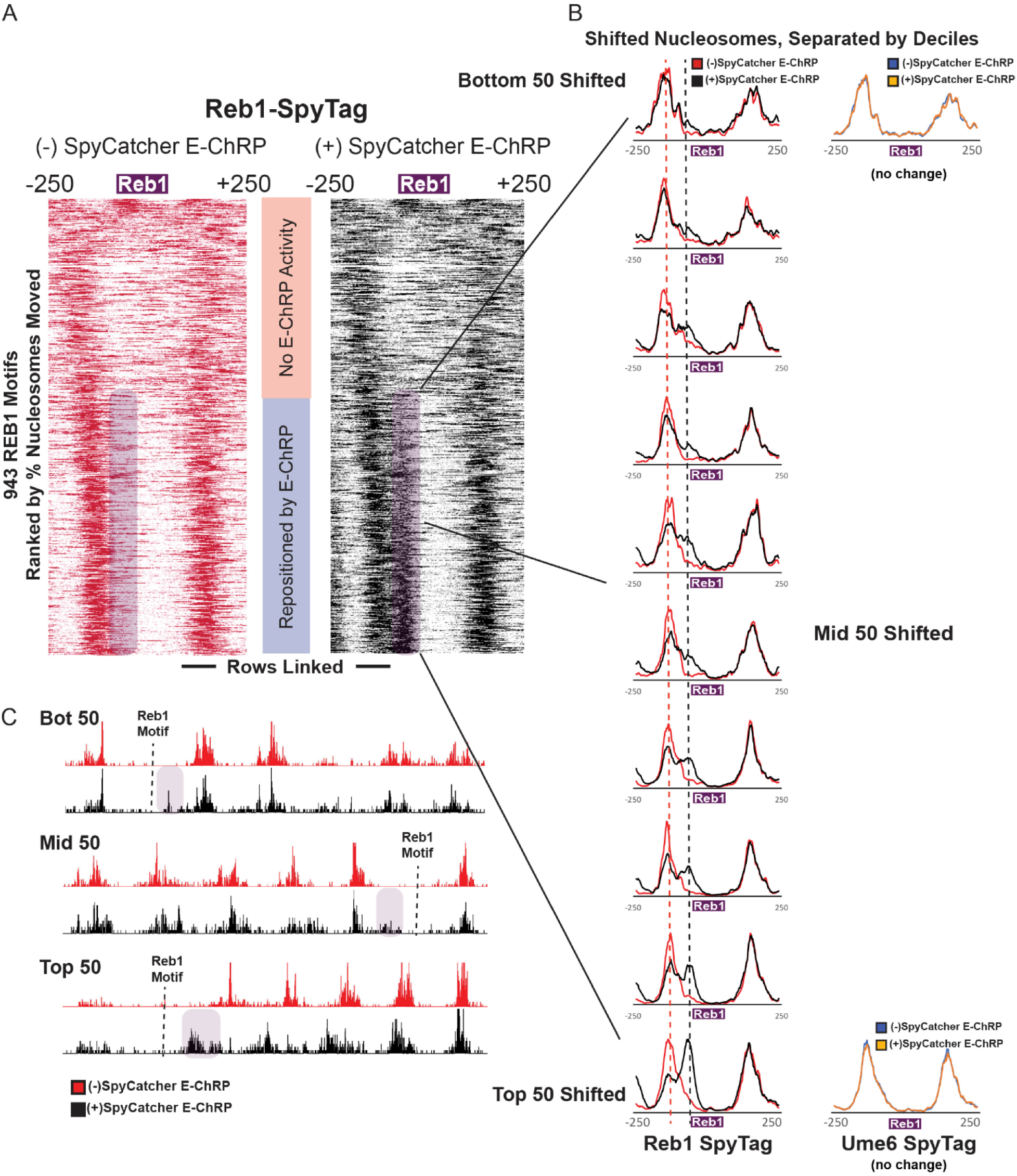
E-ChRP Targeting to Chromatin-Bound Reb1 Provides Differential Occupancy Information at Reb1 Motifs. (A) Nucleosome dyad signal at 943 intergenic Reb1 binding motifs in Reb1-SpyTag strains before (left) and after (right) 2-hour induction of SpyCatcher E-ChRP. Rows are ordered by change in nucleosome positioning after galactose induction. Purple shading highlights the region to which nucleosomes are moved by SpyCatcher E-ChRP in the Reb1-SpyTag strain. (B) The purple mobile fraction from (A) was split into deciles (∼50 motifs per decile) showing average positioning by SpyCatcher E-ChRP for each decile. Dashed lines indicate the pre-induction, unremodeled position (red) or post-induction, remodeled position (black). Ume6-SpyTag control traces are provided for the top and bottom deciles demonstrating that SpyCatcher E-ChRP cannot function at Reb1 sites in the presence of Ume6-SpyTag instead of Reb1-SpyTag. (C) Genome Browser images for representative loci showing positioning by SpyCatcher E-ChRP in a Reb1-SpyTag strain for the top, middle and bottom deciles. Purple shading indicates the motif-proximal, repositioned nucleosomes. Dashed lines indicate the location of Reb1 motif. (See also Figure S4)

To better validate the ability of SpyCatcher E-ChRP to identify fractional Reb1 occupancy at Reb1 binding sites, we compared our data set to crosslinking ChIP, CUT&RUN (Skene and Henikoff, 2017), ORGANIC (Kasinathan et al., 2014) and ChEC-Seq (Zentner et al., 2015) data sets. There was striking correlation between our data, ORGANIC and ChEC-Seq, with some motifs exclusively showing Reb1 occupancy when measured by these three methods (Fig S4). Relative Reb1 occupancies mapped by CUT&RUN and standard ChIP were less correlated with our data, suggesting that formaldehyde-free binding profiles similarly capture relative TF occupancy. Minimally, the observation that all nucleosomes are shifted at some Reb1 binding sites in a population of cells argues that some Reb1 sites are nearly 100% occupied, as E-ChRP-derived nucleosome movement cannot be observed without Reb1 binding (Fig 6C). While these Reb1 occupancy estimates are conflated with presence, accessibility and relative occupancy of motif-proximal nucleosomes, our data suggest that SpyCatcher E-ChRPs can serve as a relative measure of protein localization in cells that is orthogonal to ChIP, allowing for a lower-limit estimate of SpyTagged protein occupancy at individual binding motifs in the genome.

## dCas9-targeted Nucleosome Positioning with Nonstandard gRNAs

While the E-ChRPs described above show robust nucleosome positioning activity when targeted through various DNA binding domains or through SpyCatcher/SpyTag pairs, their ability to alter nucleosome positions depends on the interaction between pre-existing DNA binding domains with defined DNA motifs. To overcome this limitation and allow targeted positioning of single nucleosomes by design, we created a dCas9 E-ChRP (Fig 7). This construct allows versatile targeting to specific nucleosomes by designing proximal gRNAs. We recombinantly expressed the dCas9 E-ChRP in *E. coli* and purified the ∼300kDa fusion protein. To test its ability to move gRNA-targeted nucleosomes we reconstituted end-positioned mononucleosomes and designed gRNAs with or without complementarity to the extranucleosomal DNA. Successful gRNA-stimulated chromatin remodeling would result in movement of the nucleosome toward the target sequence, producing a slower-migrating centrally-positioned nucleosome (Fig 7A). While nucleosomes were efficiently moved toward the center of DNA fragments with control Chd1 protein, introduction of Chd1-dCas9 and complementary gRNA resulted in supershifted complexes with unresolved nucleosome positions. Even in the presence of 1000-fold competitor DNA for 3 days, the Chd1-dCas9 fusion protein would not release from gRNA-targeted nucleosomes (Fig 7B). This inability of dCas9 to release from target sequences is consistent with the ability of dCas9 to specifically bind and interfere with transcription in cells due to stable R-loop formation (Jinek et al., 2012; Laughery et al., 2019; Qi et al., 2013). Importantly, the inability of the dCas9 E-ChRP to release from substrate prevents its utility for precise, gRNA-targeted nucleosome positioning.

**Figure 7.**
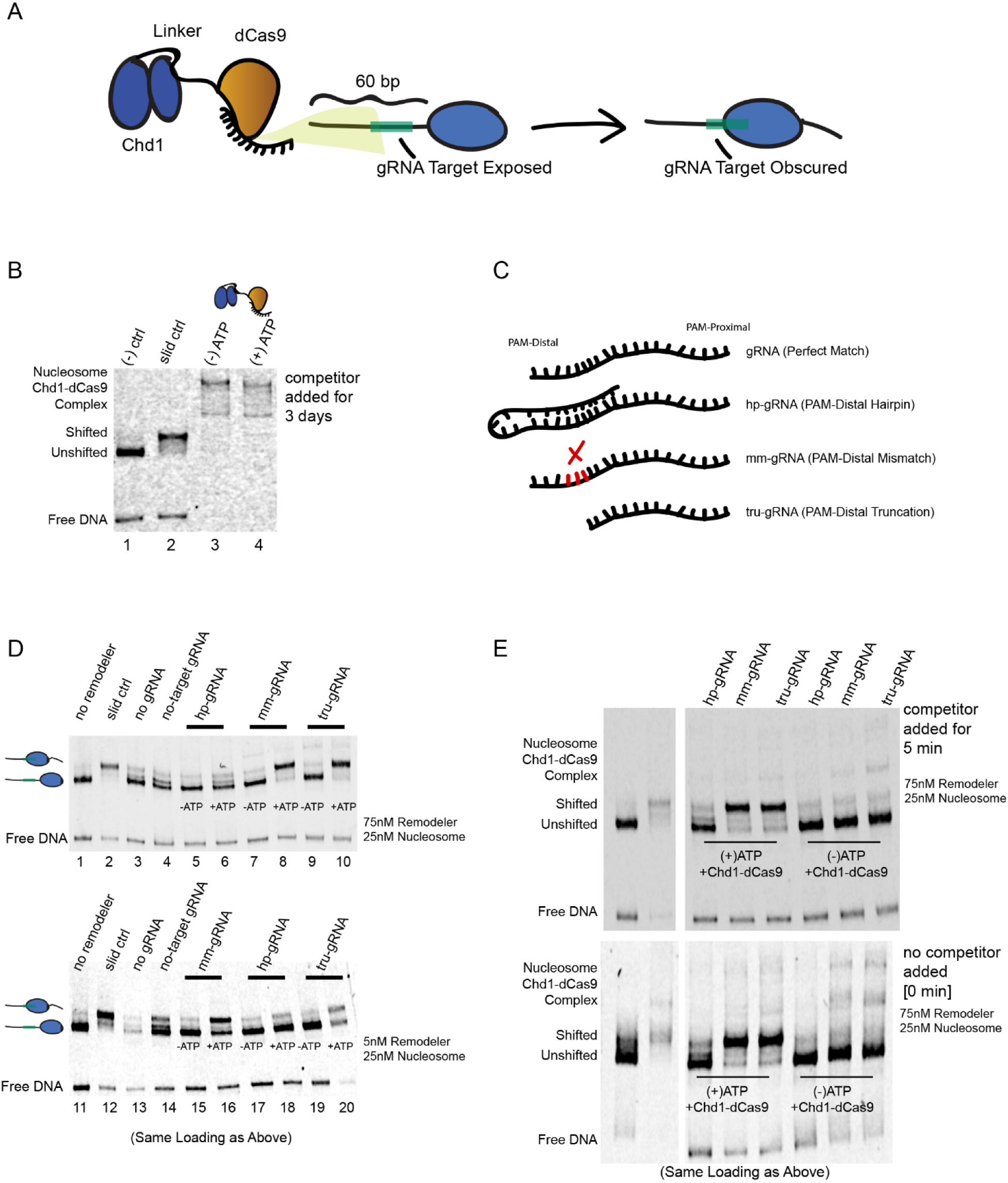
Remodeling Activity can be Targeted using a dCas9 E-ChRP with Noncanonical gRNA Substrates. (A) Cartoon depiction of dCas9-targeted chromatin remodeling and predicted repositioning of target nucleosome. (B) Nucleosome sliding assay showing irreversible association of the dCas9 E-ChRP with target nucleosomes and free DNA. The “slid ctrl” includes ATP and a control Chd1 protein capable of positioning nucleosomes toward the center of the DNA fragment. Excess unlabeled competitor DNA was added three days prior to loading. Nucleosome concentration was 25nM and E-ChRP concentration was 75nM. (C) Cartoon depiction of canonical and nonstandard gRNA protospacers. (D) Nucleosome sliding assay demonstrating robust positioning of nucleosomes by a dCas9 E-ChRP targeted with nonstandard gRNAs in single-turnover (top) or multi-turnover (bottom) conditions. The slid ctrl (lanes 2 and 12) includes ATP and a control Chd1 protein. The “no gRNA” lanes (3 and 13) contains a dCas9 E-ChRP, ATP and no gRNA. No-target gRNA samples (lanes 4 and 14) contain a dCas9 E-ChRP and a gRNA without any sequence complementarity to the substrate nucleosome. Note that the order of lanes between the upper and lower panels are similar except for loading of hp-gRNA (lanes 5,6,17,18) and mm-gRNA (lanes 7,8,15,16). (E) Nucleosome sliding assay demonstrating lack of stable association of dCas9 E-ChRPs with target nucleosomes or DNA in the presence of nonstandard gRNAs. For the upper gel, competitor DNA was added prior to loading. The bottom gel contains the identical reactions in the same order as the upper gel, but no competitor DNA was added before loading. (See also Figure S5)

To promote release of the dCas9 E-ChRP from nucleosome substrates, we used gRNAs with noncanonical structures (Fig 7C) including a truncated gRNA (Fu et al., 2014) containing only 14nt of complementarity to target sequences, a gRNA with a PAM-distal hairpin (Josephs et al., 2015) that has predicted self-annealing capacity and reduced affinity for target sequences, and a 20nt gRNA with a PAM-distal 3nt-mismatch (mm-gRNA) that would result in an R-loop with a frayed end. Both the tru-gRNA and the mm-gRNA allowed for efficient targeted repositioning of nucleosomes toward the gRNA binding site either through direct Chd1-dCas9 fusion or introduction of Chd1-SpyCatcher and dCas9-SpyTag pairs (Fig 7D, lanes 1-10 and Fig S5). These noncanonical gRNAs promoted multi-turnover catalysis by the dCas9 E-ChRP demonstrating that the weakened dCas9/gRNA complexes were stable enough to promote specific enzymatic activity but weak enough to readily and repeatedly disengage from its substrate (Fig 7D, lanes 11-20). Strikingly, dCas9-Chd1 targeted through weakened gRNAs did not require any competitor DNA to disengage from nucleosome substrates (Fig 7E). We believe this ability to readily dissociate from DNA targets while providing enough dwell time and specificity for targeted nucleosome positioning provides a facile method to alter nucleosome positions by design. Furthermore, readily-dissociating mm-gRNAs can likely be employed for dCas9-targeted epigenome editing in cells, since current epigenome editing with canonical gRNAs likely leads to a combination of local epigenetic modification and stably-bound, mutagenic R-loop formation from strong dCas9 binding (Laughery et al., 2019).

## Discussion

In conclusion, we have created and validated the use of E-ChRPs as an easy and versatile method for altering the positions of specific nucleosomes both *in vitro* and *in vivo*. We have demonstrated that E-ChRPs have widespread compatibility with various DNA binding domains and have created a single SpyCatcher E-ChRP that can be inducibly attached to chromatin-associated factors to move adjacent nucleosomes. We have shown that induced positioning of nucleosomes by E-ChRPs can establish new nucleosomal arrays, occlude transcription factor binding motifs across the genome, and report on relative transcription factor occupancy at target motifs. Finally, we have optimized a dCas9-targeted E-ChRP by creating weakened, noncanonical guide RNAs with PAM-distal mismatches to robustly position and release from targeted nucleosomes *in vitro*.

We envision future research can employ E-ChRPs to probe questions directly relating the position of nucleosomes to downstream biological processes and can lead to insight into how cells can tolerate or correct ectopic nucleosome positioning events. We further expect weakened gRNAs will be an effective strategy to target specific epigenetic changes or nucleosome positioning changes in cells while limiting indirect consequences of stably- or irreversibly-bound dCas9 (Laughery et al., 2019). Finally, the ability to position nucleosomes onto target sequences may potentially lead to the development of E-ChRPs that block oncogenic or other disease-related transcription factors from accessing binding sites genome-wide.

## Limitations

While the E-ChRPs described in this work are highly versatile allowing for multiple targeting schemes, there are some limitations in the ability of E-ChRPs to position target nucleosomes. First, to create a functional Chd1-TF(DBD) fusion, the boundary of the DNA binding domain for the specific transcription factor must be known, and it must fold in the context of the fusion protein. While all fusions we have tested have been functional to date, we focused on well-behaved and well-studied DNA binding domains. Second, there is still a limitation on how far E-ChRPs can “reach”. Based on our *in vivo* mapping results, if a nucleosome edge is initially beyond ∼75 base pairs from the E-ChRP recruitment site, nucleosome repositioning activity is less favorable. Moreover, E-ChRPs do not appear to have any *de novo* nucleosome deposition activity, so exaggerated nucleosome-free regions of the genome would not permit nucleosome positioning. While we are very interested in the ability of SpyCatcher E-ChRPs to position different fractions of nucleosomes at different sites in the genome and we speculate this is due to relative SpyTagged TF occupancy, our method is blind to TF binding sites where there are no motif-proximal nucleosomes. Further, it would be challenging to use E-ChRPs in isolation to define TF binding landscape due to the relatively noisy signal generated from fractional repositioned nucleosomes. This is especially true for sites where small fractions of nucleosomes are repositioned. Finally, while the determination that dCas9 E-ChRPs work readily with mismatched gRNAs may allow better assessment of targeted chromatin modification effects (since dCas9 does not remain stably associated with the target sequence), we note that mismatched gRNAs naturally possess a lower capacity for target specificity. Nevertheless, the robust activity we see with dCas9 E-ChRPs and mismatched gRNAs may be generally useful for assessing metastable off-target locations of gRNAs.

## Methods

### Plasmids, Strains, and Growth Conditions

The initial Ume6 E-ChRP scaffold was prepared by Gibson assembly (Gibson et al., 2009) in p416-ADH1 (Mumberg et al., 1995) to include an N-terminal NLS (KKKRK), residues 118-1000 of *S. cerevisiae* Chd1, nine repeats of glycine-glycine-serine, residues 1001-1014 of *S. cerevisiae* Chd1, two additional repeats of glycine-glycine-serine, an AfeI restriction site, residues 764-836 from S. cerevisiae Ume6 and a HindIII restriction site. An analogous backbone was also created with residues 118-1000 of *S. cerevisiae* Chd1 linked directly to the two glycine-glycine-serine repeats followed by AfeI, Ume6 DBD, and HindIII site. Cloning vectors were created in pDEST17, p416-TEF, p416-GPD, p426-GPD and HO-pGAL-poly-KanMX4-HO (Voth et al., 2001) (a gift from David Stillman, Addgene 51664) to swap the C-terminal Ume6 domain with other targeting domains using AfeI/HindIII restriction cloning, sticky-end PCR or Gibson Assembly (Gibson et al., 2009; Walker et al., 2008). Fusions used in this study include *S. cerevisiae* Ume6 (residues 764-836, cloned from yeast genomic DNA), *D. melanogaster* Engrailed (residues 454-543, cloned from fly genomic DNA), *S. pombe* Res1 (residues 1-147, cloned from a gBlock), *R. norvegicus* Glucocorticoid Receptor (residues 428-513, cloned from a gBlock), *E. coli* AraC (residues 175-281, provided by Gregory Bowman), the SpyCatcher domain (Zakeri et al., 2012) (a gift from Mark Howarth, Addgene 35044) and dCas9 (subcloned from Addgene 49013, a gift from Timothy Lu (Farzadfard et al., 2013)). Non-integrating plasmids (p416- or p426-) were transformed into *S. cerevisiae* strain W303 (RAD5+) and grown in SD-Ura overnight, diluted to OD_600_=0.2 in SD-Ura and grown to OD_600_=0.6-0.8 for chromatin analysis. For galactose induction of E-ChRPs, cells were grown in YP media with 2% raffinose as the sole carbon source. In mid-log phase, raffinose (− induction) or galactose (+ induction) was added to a final concentration of 2% and cells were grown for 2 additional hours at 30°C with shaking. Cells were then fixed and harvested for chromatin analysis. To make SpyTagged yeast strains, a C-terminal 3x-FLAG tag followed by the SpyTag sequence (AHIVMVDAYKPTK) was added by integration at the endogenous locus for the protein of interest using a selectable drug marker. To recombinantly express SpyTagged DNA binding domains for biochemical analysis, domains of interest were amplified by PCR from source (yeast or fly genomic DNA) followed by restriction cloning into a pDEST-MCS-SpyTag vector.

### Protein Purification

Chd1 constructs were expressed from pDEST17 vectors (Invitrogen) as previously described (Hauk et al., 2010). Briefly, proteins were expressed in BL21(DE3) cells, with the RIL plasmid (Stratagene) to aid expression and a plasmid expressing the Trigger Factor chaperone (a kind gift from Li Ma and Guy Montelione) to improve solubility. After induction with 300uM isopropyl-D-thiogalactopyranoside (IPTG) and growth at 18°C for 16 h, cells were lysed by sonication in 500 mM NaCl, 10% glycerol, and 25 mM Tris, pH 7.8. Lysate was clarified by centrifugation at 25,000xg and soluble protein was purified using Co^2+^ affinity chromatography (TALON column, GE Healthcare) followed by anion-exchange chromatography (Q-FF, GE Healthcare).

### Nucleosome Sliding Assay

Recombinant yeast histones were purified as previously described (Luger et al., 1999) and dialyzed by gradient salt dialysis onto the Widom 601 positioning sequence (Lowary and Widom, 1998). Nucleosome sliding was performed at 25°C in sliding buffer (50 mM KCl, 15 mM HEPES, pH 7.8, 10 mM MgCl_2_, 0.1 mM EDTA, 5% sucrose, 0.2 mg/ml bovine serum albumin [BSA], with or without 5 mM ATP) by incubating 0, 10, 20, 40, or 60nM purified E-ChRP with 30nM reconstituted mononucleosomes for 100 minutes. SpyCatcher reactions included 25, 50, or 100nM Chd1-SpyCatcher and 0, 50 or 100nM of the SpyTagged DNA binding domain with 30nM of labeled mononucleosomes. Unless specifically indicated, mononucleosomes contained E-ChRP recognition sequences in the extranucleosomal DNA. Reactions were quenched by diluting 1:2 with solution containing 3 uM competitor DNA (with E-ChRP recognition sequences) and 5% sucrose. Native PAGE (6%) was used to separate the positioning of the mononucleosomes, with Cy5.5-labeled nucleosomal DNA detected by a LiCor Odyssey FC imager. Chd1-dCas9 experiments were performed with indicated concentrations of nucleosomes and remodeler in the presence or absence of 5mM ATP for 100 minutes. Guide RNAs were synthesized from a T7 promoter using the SureGuide gRNA Synthesis Kit (Agilent). gRNA was added in a 2-fold excess relative to dCas9 concentration. For Chd1-SpyCatcher-SpyTag-dCas9 reactions (Figure S5), 10-fold excess, concentrated Chd1-SpyCatcher was pre-incubated with SpyTag-dCas9 for 1 hour prior to dilution and reaction initiation.

### Micrococcal Nuclease Digestions and Library Construction

Micrococcal nuclease digestions were performed as previously described (Rodriguez et al., 2014). Briefly, cells were grown to mid-log phase and fixed with 1% formaldehyde. Chromatin was digested with 10, 20, and 40 units of MNase for 10 minutes. Proper nuclease digestion of DNA was analyzed by agarose gel and samples with approximately 80% mononucleosomes were selected for library construction. After crosslink reversal, RNase treatment, Calf Intestine Phosphatase (CIP, NEB) treatment and Proteinase K digestion, mononucleosome-sized fragments were gel-purified and used to construct libraries with the NuGEN Ovation Ultralow kit per the manufacturer’s instructions. Libraries were sequenced at the University of Oregon’s Genomics and Cell Characterization Core Facility on an Illumina NextSeq 500 on the 37 cycle, paired-end, High Output setting, yielding approximately 20 million paired reads per sample.

### Chromatin Immunoprecipitation and Library Construction

Chromatin immunoprecipitation was performed as previously described (Rodriguez et al., 2014). Briefly, cells were grown to mid-log phase, fixed with 1% formaldehyde, and lysed by bead-beating in the presence of protease inhibitors. Chromatin was fragmented by shearing in a Bioruptor sonicator (Diagenode) for a total of 30 minutes (high output, 3×10’ cycles of 30 sec. on, 30 sec. off). Sonication conditions were optimized to produce an average fragment size of ∼300 basepairs. FLAG-tagged protein was immunoprecipitated using FLAG antibody (Sigma) and Protein G magnetic beads (Invitrogen). After crosslink reversal and Proteinase K digestion, DNA was purified using Qiagen MinElute columns and quantified by Qubit High-Sensitivity fluorometric assay. Libraries were prepared using the NuGEN Ovation Ultralow kit by the manufacturer’s instructions and sequenced at the University of Oregon’s Genomics and Cell Characterization Core Facility on an Illumina HiSeq4000 with 50 or 100 cycles of single-end setting, yielding approximately 15 million reads per sample.

### Data Processing and Analysis

MNase sequencing data were analyzed as described previously (McKnight et al., 2015; McKnight and Tsukiyama, 2015; McKnight et al., 2016). Briefly, paired-end reads were aligned to the *S. cerevisiae* reference genome (Cunningham et al., 2015) with Bowtie 2 (Langmead and Salzberg, 2012), and filtered computationally for unique fragments between 100 and 200 bp. Dyad positions were calculated as the midpoint of paired reads, then dyad coverage was normalized across the *S. cerevisiae* genome for an average read/bp of 1.0. Nucleosome alignments to transcription factor binding sites were performed by taking average dyad signal at each position relative to all intergenic instances of a motif center. Motifs were obtained from the JASPAR database (Khan et al., 2018) and intergenic instances were found using the Saccharomyces Genome Database Pattern Matching tool (http://www.yeastgenome.org/cgi-bin/PATMATCH/nph-patmatch). Specifically, the Reb1 motif was defined as TTACCC(G/T) and Ume6 motif was WNGGCGGCWW. For ChIP-seq data, single-end reads were aligned to the *S. cerevisiae* reference genome with Bowtie 2 and total read coverage was normalized such that the average read at a genomic location was 1.0. ChIP peaks were called using a 400 bp sliding window with a threshold average enrichment within the window of 4.0. Reb1 ORGANIC data (Kasinathan et al., 2014) were from the “80mM IP” sample (SRX263794); Reb1 CUT&RUN data (Skene and Henikoff, 2017) were from merged “cut-and-run 8s” and “cut-and-run 16s” samples (SRX2009989 and SRX2009990); Reb1 ChEC-seq data (Zentner et al., 2015) were from the “Reb1 ChEC-seq 30s” (SRX974362). Data were visualized using Integrated Genome Browser (Freese et al., 2016).

## Author Contributions

Conceptualization, D.A.D., L.E.M., and J.N.M.; Methodology, D.A.D., J.G.C., O.G.B.B., Z.D.J., L.E.M., and J.N.M.; Investigation, D.A.D., J.G.C., O.G.B.B., Z.D.J., L.E.M., and J.N.M., Writing - Original Draft, D.A.D., J.G.C., and J.N.M.; Writing - Review & Editing, D.A.D., J.G.C., Z.D.J., L.E.M., and J.N.M.; Visualization, D.A.D. and J.N.M.; Supervision, L.E.M. and J.N.M.; Project Administration, J.N.M.; Funding Acquisition, J.N.M.

## Acknowledgements

We would like to thank the Bowman lab (Johns Hopkins University) for histone plasmids and Chd1 constructs, the Tsukiyama lab (Fred Hutchinson Cancer Research Center) and Zentner lab (Indiana University) for selectable epitope tagging vectors and the Doe Lab (University of Oregon) for a fly genomic DNA extraction protocol. This work was supported by a National Institutes of Health training grant T32 GM007759 (to D.A.D. and O.G.B.B.), by a National Institutes of Health training grant T32 GM007413 (to D.A.D.) and by NIGMS R01 GM129242 (J.N.M.) and the Donald and Delia Baxter Foundation (J.N.M.).

## Supplemental Figures and Legends

**Figure S1, Related to Figure 3.**
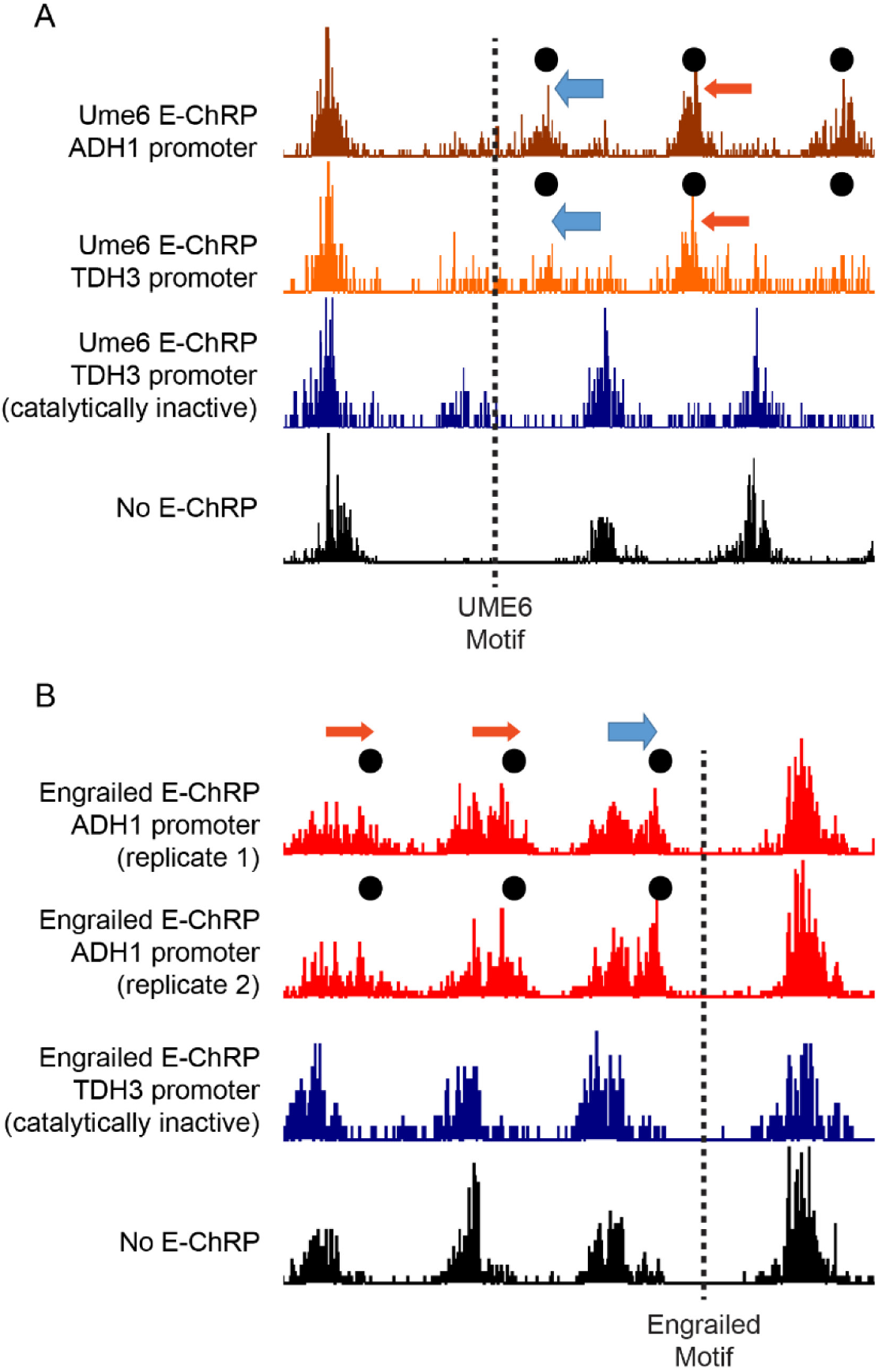
Nucleosome Positioning by E-ChRPs Requires Chd1 Catalytic Activity. (A) Representative locus showing changes in nucleosome signal after lower (dark red) or higher (orange) expression levels of catalytically-active Ume6 E-ChRP. Black circles mark altered nucleosome positions compared to a catalytically-inactive E-ChRP with a Walker B D513N mutation in the Chd1 remodeling core (blue) or a parental strain (black). (B) Representative locus showing altered nucleosome positions at an Engrailed binding motif by catalytically-active (red) but not catalytically-inactive (blue) Engrailed E-ChRP compared to a parent strain (black). Respective motif locations are indicated with a dashed line.

**Figure S2, Related to Figure 4.**
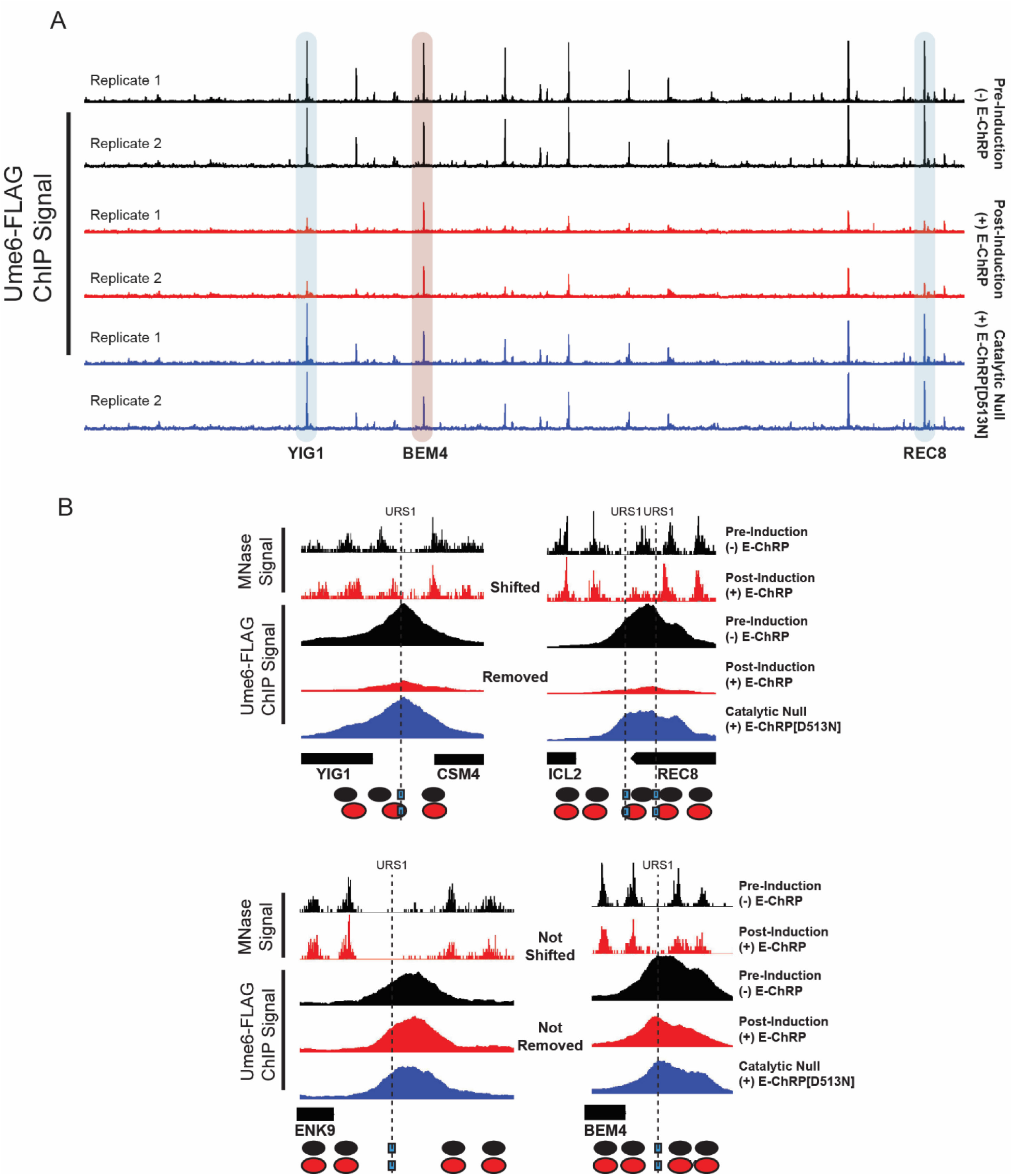
Eviction of Ume6-FLAG by a Catalytically-Active E-ChRP. (A) Genome Browser image showing biological replicates of Ume6-FLAG ChIP signal across ChrXVI before (black) and after (red) 2-hour induction of Ume6 E-ChRP with galactose. Blue peaks correspond to a Ume6-FLAG ChIP signal in the presence of a catalytically inactive Ume6 E-ChRP containing a Walker B (D513N) substitution in the Chd1 remodeling core. Blue and red highlights correspond to loci in (B). (B) Representative examples of E-ChRP targets where endogenous Ume6 is removed (top) or not removed (bottom) after galactose induction of Ume6 E-ChRP. The catalytic null E-ChRP retains a Ume6 DBD but has a Walker B (D513N) substitution in the Chd1 catalytic core.

**Figure S3, Related to Figure 5.**
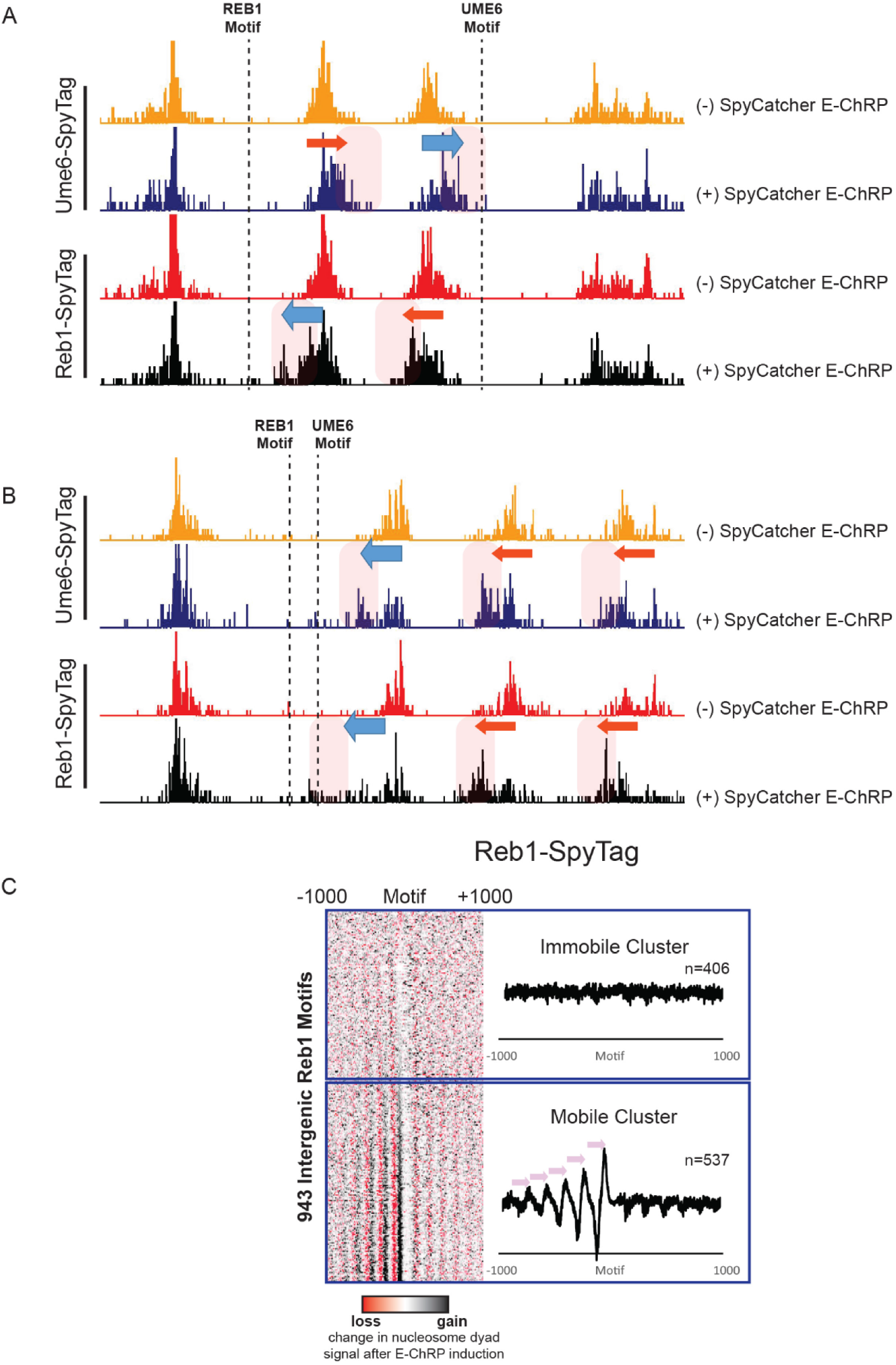
Targeted Chromatin Remodeling with Chd1-SpyCatcher and SpyTagged Chromatin Factors. (A) Genome Browser image showing nucleosome positions before and after induction of SpyCatcher E-ChRP in cells containing Ume6-SpyTag (top) or Reb1-SpyTag (bottom). Location of a proximal Reb1 binding motif or Ume6 binding motif is denoted by a dashed line while directional nucleosome positioning is indicated by blue and red arrows. At this locus, Reb1-SpyTag and Ume6-SpyTag cause different nucleosomes to be selectively moved by SpyCatcher E-ChRP as directed by the location of bound Ume6 or Reb1. (B) Same as (A) showing a locus where Reb1 and Ume6 motifs are adjacent to each other. In this case, both Reb1-SpyTag and Ume6-SpyTag allow the SpyCatcher E-ChRP to select the same nucleosome but the nucleosome is moved to different final locations based on the location of the individual bound factors. Reb1-SpyTag leads to further positioning than Ume6-SpyTag because the Reb1 motif is distal to the Ume6 motif. (C) Heat map (left) showing the difference in nucleosome dyad signal +/− 1000 bp from 943 Reb1 motifs after SpyCatcher E-ChRP induction in Reb1-SpyTag cells. Average change in nucleosome signal after SpyCatcher E-ChRP induction at Reb1 motifs where nucleosomes are moved (mobile cluster) or not moved (immobile cluster) are provided (right).

**Figure S4, Related to Figure 6.**
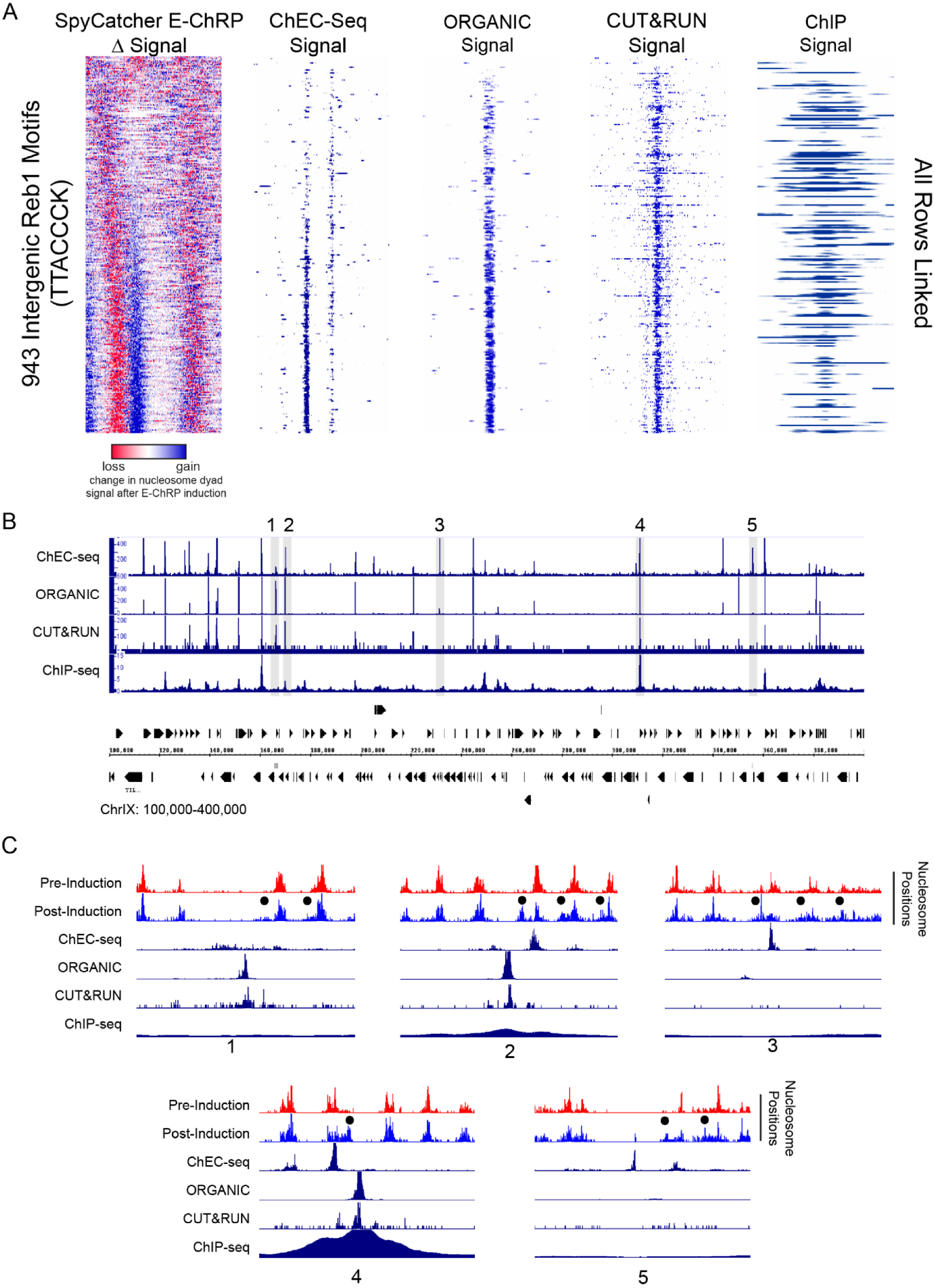
Comparison of SpyCatcher-Induced Chromatin Remodeling at Reb1-SpyTag Sites to Previous Reb1 Mapping Strategies. (A) Analysis of Reb1 binding at 943 intergenic Reb1 motifs (TTACCCK) using indicated methods for Reb1 mapping. All data are ordered based on the ranked change in nucleosome positioning after SpyCatcher E-ChRP induction in a Reb1-SpyTag strain (left). All data are centered at the Reb1 motif and display +/− 250 base pairs from each motif. (B) Genome Browser image showing Reb1 binding across ChrIX for indicated Reb1 mapping strategies. Highlighted regions of interest are displayed in (C). (C) Zoomed-in Genome Browser images showing nucleosome repositioning by SpyCatcher E-ChRP at Reb1-SpyTag sites (blue versus red) and relative Reb1 signal from indicated methods. All regions show nucleosome shifts by SpyCatcher E-ChRP (black circles), ChEC-seq signal and ORGANIC signal. Regions 2 and 4 show Reb1 binding using all methods. Regions 1,3 and 5 lack ChIP signal. Regions 3 and 5 lack CUT&RUN signal. Regions 3 and 5 have very low but detectable ORGANIC signal despite significant nucleosome shifts by SpyCatcher E-ChRP and high ChEC-seq signal. Numbering corresponds to (B).

**Figure S5, Related to Figure 7.**
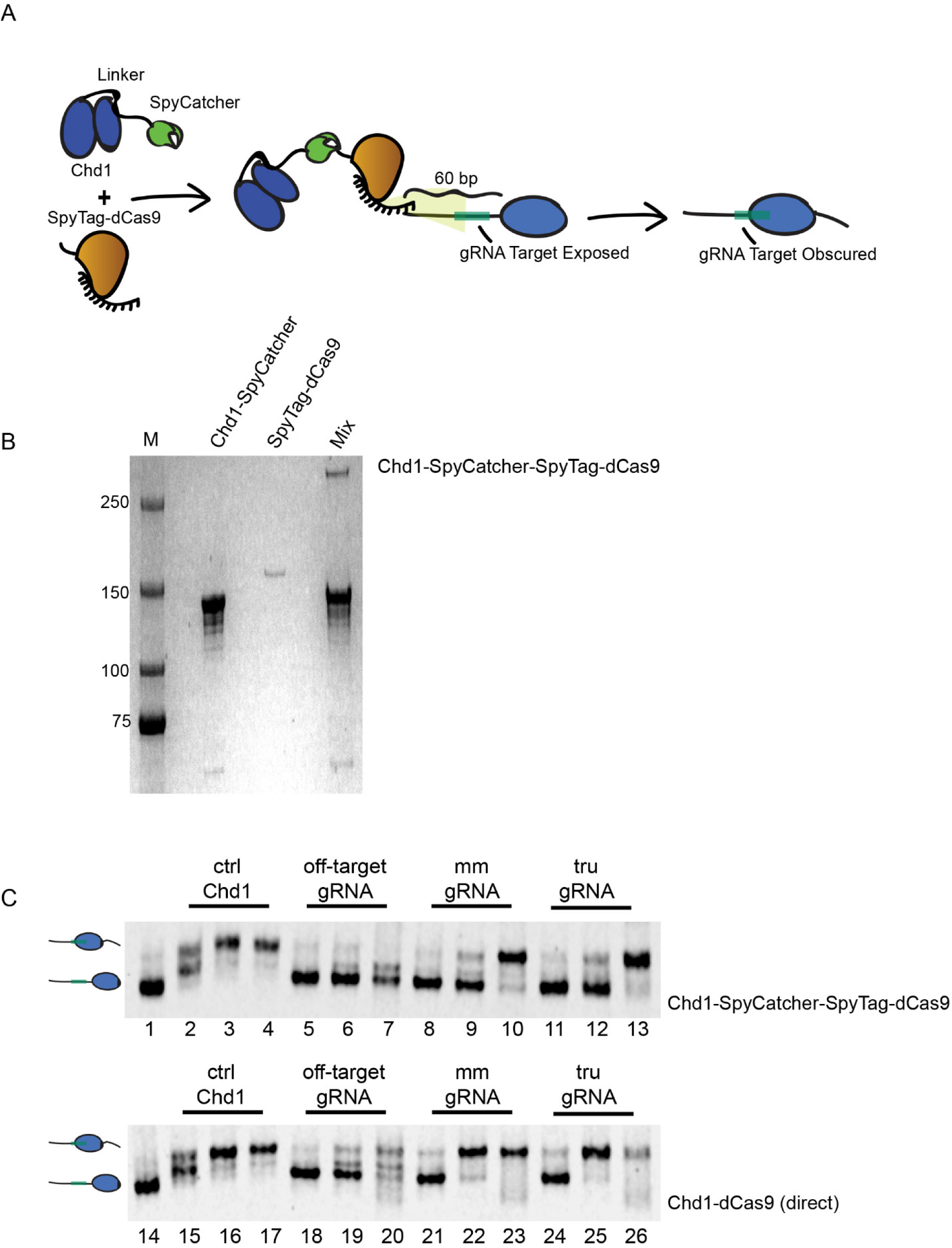
Noncanonical gRNAs Stimulate E-ChRP Activity for a Split Chd1-SpyCatcher and SpyTag-dCas9 Pair. (A) Cartoon representation of Chd1-SpyCatcher combining with SpyTag-dCas9 to form a functional, gRNA-targeted E-ChRP and predicted nucleosome positioning at a target nucleosome. (B) SDS-PAGE demonstrating full conversion of SpyTag-dCas9 to Chd1-SpyCatcher-SpyTag-dCas9 in the presence of excess Chd1-SpyCatcher prior to remodeling assays. (C) Comparison of Chd1-SpyCatcher-SpyTag-dCas9 remodeling activity (top) on target nucleosomes to Chd1-dCas9 (direct fusion) activity (bottom) using indicated gRNAs. A catalytically active Chd1-Ume6 protein was used as a positive control (ctrl Chd1) for nucleosome positioning. Lanes 1 and 14 contain unremodeled nucleosome (25nM). Lanes 2-4 and 15-17 include 1.5, 15 and 150nM Chd1-Ume6. All other lanes contain the indicated Chd1-dCas9 (either direct fusion or SpyCatcher/SpyTag pair) with 1.5, 15 and 150nM remodeler for each gRNA condition. For “off-target gRNA” conditions, a gRNA with no complementarity to the nucleosome substrate was included in the reaction.

